# Adventitial leptin receptor-expressing fibroblasts are preferential contributors to fibrotic remodeling of the heart post infarction

**DOI:** 10.1101/2025.10.14.682477

**Authors:** Veronica Larcher, Ariane Fischer, Lunfeng Zhang, Omar Almolla, Mattia Chiesa, Francesca Andriani, Chiara Wernet, Simone Serio, Lukas Tombor, Sofia Peruzzo, Debanjan Mukherjee, Rossana Bussani, Serena Zacchigna, Ralf H. Adams, Stefanie Dimmeler, Sylvia M. Evans, Paola Cattaneo, Nuno Guimarães-Camboa

## Abstract

**Background:** Cardiac fibrosis, a hallmark of heart failure and an unmet clinical need, arises from pathological activation of pre-existing cardiac fibroblasts (CFs), <u>but the contribution of CF heterogeneity to this process remains unclear.</u>

**Methods:** Murine models were used to lineage trace or deplete a specific sub-population of CFs at baseline and after myocardial infarction (MI). Transcriptional and epigenetic differences between fibroblast subsets were assessed using next-generation sequencing. Conservation in humans was evaluated through single-cell RNA-seq datasets and histological examination.

**Results:** In mice, fibroblasts were the sole cardiac cell type expressing the signaling-capable isoform of the leptin receptor (LepR). LepR+ CFs emerged neonatally, occupied a defined niche in the coronary adventitia, exhibited enhanced hedgehog signaling, <u>and responded to leptin</u>. <u>After MI, LepR-Cre+ CFs proliferated more than interstitial CFs, became a predominant fibroblast lineage in the scar, and their genetic ablation reduced fibrosis while improving function. LepR+ CFs were also detected in the human heart, where they were embedded in an adipocyte-rich niche.</u>

**Conclusions:** These findings identify adventitial fibroblasts as key drivers of pathological remodeling and demonstrate that fibroblasts, rather than cardiomyocytes, are the principal responders to leptin in the heart, redefining how this major endocrine pathway influences cardiac remodeling and disease.

## INTRODUCTION

Despite advances in clinical care, heart failure remains a leading cause of mortality worldwide^1^. Cardiac fibrosis, resulting from excessive deposition of extracellular matrix (ECM) by activated cardiac fibroblasts (CFs), is a hallmark of most types of heart failure and an unmet clinical need^2,3^. Several studies have shown that most activated CFs arise from pre-existing quiescent CFs^4–7^, however, it remains unclear whether all pre-existing CFs contribute equally to fibrotic processes. In the context of hypertension, fibrosis often develops in the vicinity of large vessels (perivascular fibrosis)^4^. Recent efforts to target fibrosis with CAR T-cells have shown efficient elimination of interstitial fibroblasts, while sparing fibroblasts located in the adventitia of large coronary vessels^8^. These observations suggest that CFs from the adventitia of large coronary vessels might possess distinct molecular features that enable them to play a pivotal role in pathological remodeling. Yet, this hypothesis remains unproven, largely due to the lack of tools that allow selective labeling and tracing of specific CF subsets.

Obesity is widely recognized as a major risk factor for heart failure^9^. Beyond serving as an energy reservoir, adipose tissue is a key endocrine organ, releasing a broad range of adipokines with systemic effects^10,11^. Among these is leptin, a 16-KDa hormone secreted by adipocytes and primarily recognized by its capacity to act on hypothalamic leptin receptor (LepR)-expressing neurons, controlling food intake and energy expenditure^11^. Alternative *Lepr* splicing generates at least six distinct isoforms^12^, but only the long isoform (*Leprb*) is capable of intracellular signaling, primarily via JAK-STAT^11,12^. In the cardiovascular system, leptin has been implicated in promoting cardiac hypertrophy and regulating vascular tone and inflammation^10^. Early studies showed that leptin can induce hypertrophy of cultured rat^13^ or human^14^ ventricular cardiomyocytes. Subsequent cardiomyocyte-specific LepR ablation studies in mice suggested that leptin is essential for cardiomyocyte energy homeostasis^15,16^. The role of leptin in non-cardiomyocyte populations has been less explored, but leptin has been linked to cardiac fibrosis^17^. Studies using cultured CFs suggested this is mainly explained by direct regulation of genes encoding ECM molecules and matrix-degrading proteases^17^. However, whether these in vitro findings accurately reflect in vivo processes, and whether all CF subsets possess the molecular machinery necessary to sense leptin remains largely unknown.

We found that, contrary to previous reports, LepR expression in the murine heart is restricted to fibroblasts and identified a previously unrecognized lineage of LepR-Cre+ CFs that occupy a well-defined niche in the adventitia of large coronary arteries. LepR expression in this lineage emerged in the neonatal period, coinciding with the onset of white adipogenesis and a peak in circulating leptin. <u>Using both constitutive and inducible LepR-Cre models, we showed that following myocardial infarction (MI), LepR-Cre+ adventitial CFs proliferated more than interstitial CFs and became the dominant CF population within the scar. Ablation of LepR-Cre+ adventitial CFs prior to MI reduced fibrosis and improved cardiac function, and LepR+ adventitial CFs were also detected in human hearts. Together, these findings challenge current models of leptin signaling in the heart and define a novel lineage of fibroblasts that act as preferential progenitors during post-MI remodeling, highlighting them as a potential therapeutic target for limiting pathological fibrosis.</u>

## METHODS

### Animal Models

Procedures involving mouse models conformed to the guidelines from Directive 2010/63/EU of the European Parliament on the protection of animals used for scientific purposes or the NIH Guide for the Care and Use of Laboratory Animals. Experiments were authorized by the Regierungspräsidium Darmstadt, Hessen, Germany (FU/2041) or the UCSD IACUC. The following alleles were used: LepR-Cre^18^, Col1a1-GFP^19^, Tie2-Cre^20^, Gli1-CreERT2^21^, Rosa-tdTomato^22^, Rosa-iDTR^23^, LepR-CreERT2^24^, and Leptin-LacZ (KOMP). <u>Male and female animals were used in all experiments</u>. Detailed methods are described in the Supplemental Material.

### Human subjects

Analyses involving human samples followed the principles outlined in the Declaration of Helsinki. Portions of the left ventricular free wall containing large segments of the LAD were dissected post-mortem and immediately placed in 4% PFA for overnight fixation, followed by routine protocols for paraffin histology. Detailed protocols and subject information are provided in the Supplemental Material.

### Isolation of murine CFs

CFs were isolated from adult and neonatal murine hearts using a modified version of a previously described dissociation protocol^4^. A full description of the procedure is provided in the Supplemental Material.

### RNA-seq

FACS-sorted CFs were collected in Qiazol (Qiagen 79306) and mRNA was purified using the miRNeasy Micro Kit (Qiagen 217084) with on-column DNase digestion. Quality control was done using TapeStation (Agilent). Samples with RNA integrity number (RIN) between 6.7-9.4 were used for library preparation using the TruSeq Stranded mRNA library Prep kit (Illumina 20020594) and sequenced (single read 50 protocol) on an Illumina HiSeq 4000 System.

### ATAC-seq

Per replicate, 75000 CFs were lysed in ice-cold lysis buffer (10 mM TrisHCl pH 7.4, 10 mM MgCl2, 0.1% Igepal CA-630) and stored at –80°C in storing buffer (10 mM Hepes pH 7.5, 2 mM MgCl2, 25 mM KCl, 250 mM sucrose, 1 mM DTT, 1 mM PMSF, 70% glycerol) until further processing. Libraries were prepared using the ActiveMotif ATAC-seq kit (53150).

### Bioinformatic Analyses

Next generation sequencing datasets were analyzed in R (4.2.1) using standard pipelines. Unless otherwise stated, significance was defined as an absolute log2FC (|log2FC|) ≥ 0.5 and an adjusted p-value ≤ 0.05. Detailed procedures are available in the Supplemental Material.

### Quantitative and statistical analyses

Cell counts and RNAscope puncta were quantified using ImageJ. Statistical analyses were performd in GraphPad Prism. Data distributions were first tested for normality (Shapiro–Wilk test) and homogeneity of variance to determine the suitability of parametric versus non-parametric analysis.

### Data availability

Sequencing datasets were deposited in Zenodo and GEO (GSE253551).

## RESULTS

### A novel lineage of adventitial LepR-Cre+ Cardiac Fibroblasts

Leptin, adiponectin and resistin are adipocyte-specific cytokines. To assess whether any of these can directly signal to the heart, we evaluated expression of their cognate receptors in a cell type-resolved cardiac RNA-seq dataset^25^. Receptors for all three adipokines were detectable in the heart. Whereas receptors for adiponectin and resistin were ubiquitous, the receptor for leptin (*Lepr*) was selectively expressed by cardiac fibroblasts (CFs) (Figure 1A). To validate this highly-specific expression pattern, we performed lineage tracing studies employing a LepR-Cre that uses an IRES sequence to drive Cre expression under the control of the long, signaling-capable, isoform of LepR without disrupting the endogenous transcript^18^. By combining this allele with two reporter strains (*Rosa26-tdTomato*^22^ and *Col1a1-GFP*^4,19^), we generated triple transgenic mice in which all cells with a history of LepR expression were permanently marked by tdTomato, while all fibroblasts were labeled by GFP. Flow cytometry confirmed that expression of the long isoform of LepR was confined to CFs, with more than 99% of tdTomato+ cells co-expressing Col1a1-GFP (Figure 1B,C). Histological analyses revealed that LepR-Cre+_CFs were not uniformly distributed throughout the heart, but instead accumulated in the adventitia of large coronary vessels<u>, that extended from the ventricular base to near apical regions</u> (Figure 1D). Confocal microscopy (63x magnification) revealed that, in images containing large coronary vessels, LepR-Cre+_CFs constituted 64% of all CFs and accounted for 100% of CFs located within the immediate adventitial niche. By contrast, only 8% of interstitial CFs were LepR-Cre+ (Figure 1E). Immunostaining with specific markers revealed that LepR-Cre+_CFs localized preferentially around large arteries, identified by the presence of αSMA+ smooth muscle cells, rather than around veins, marked by Endomucin (Figure 1F).

**Figure 1.**
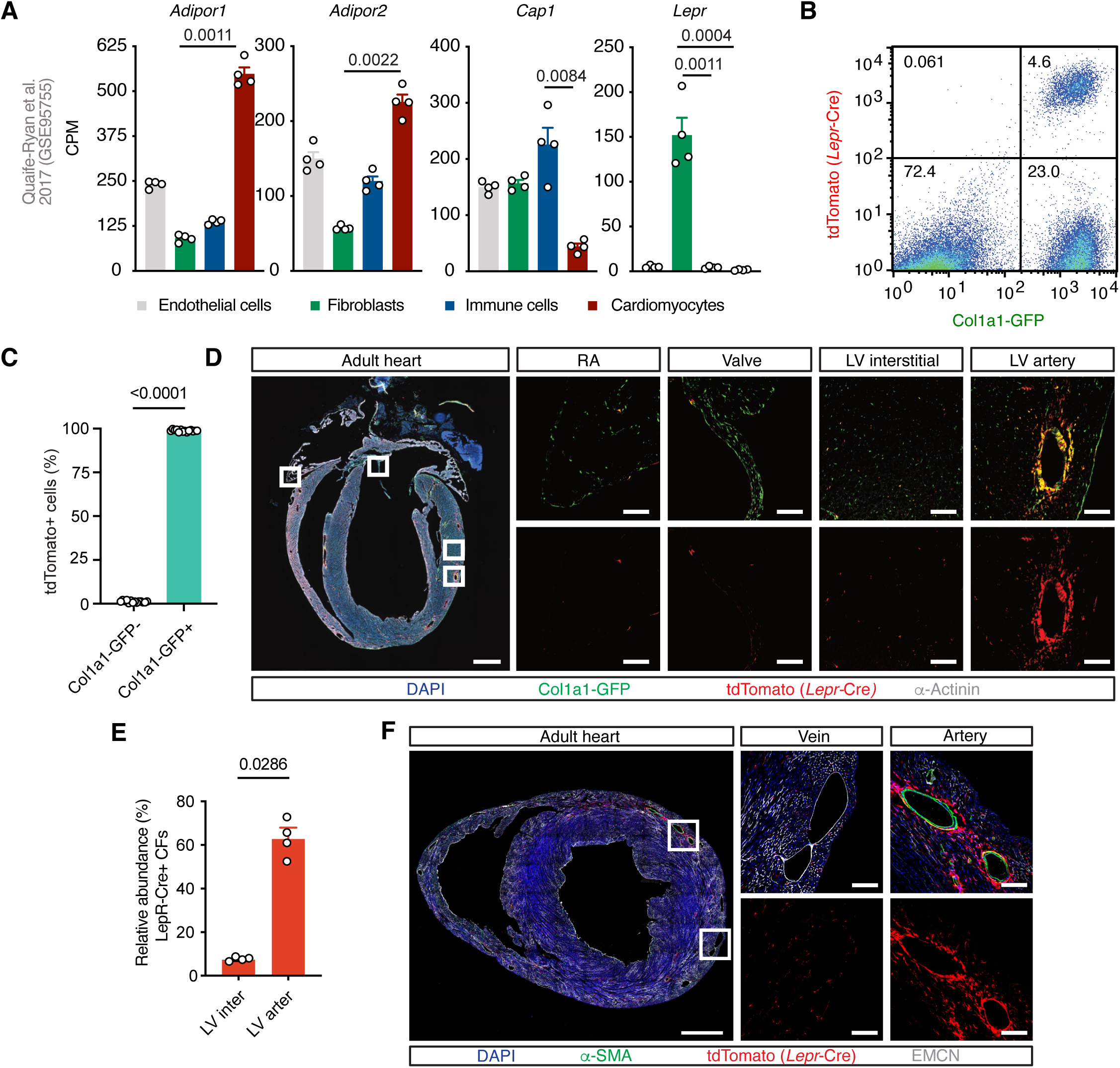
Identification of a lineage of LepR-Cre+ cardiac fibroblasts occupying the adventitia of large coronary arteries. (**A)** Expression of adipokine receptors in RNA-seq of isolated cardiac cell types. Whereas most receptors were broadly expressed, *Lepr* was selectively expressed in fibroblasts (*N* = 4, mean ± SEM, Kruskal-Wallis test). (**B,C**) Representative flow cytometry (B) and corresponding quantification (C) of cells from *Lepr-Cre/WT; Rosa26-tdTomato/WT; Col1a1-GFP*+ hearts showing that the vast majority of cells with a history of expression of the long LepR isoform (tdTomato+) were fibroblasts (Col1a1-GFP+) (*N* = 14, mean ± SEM, Mann-Whitney test). (**D**) Histological analysis and (**E**) corresponding quantification (*N* = 4, mean ± SEM, Mann-Whitney test) revealing that LepR-Cre+ CFs (GFP+, tdTomato+, yellow/orange) accumulated in the adventitia of large coronary vessels. Boxed areas are shown at higher magnification. (**E**) Immunofluorescence for Endomucin (grey, marker of endocardium and venous endothelium) and αSMA (green, marking smooth muscle cells in coronary arteries) showing LepR-Cre+ CFs localized specifically around large arteries. Scale bars: 1 mm (whole-heart images) and 100 µm (magnified panels).

Together, these findings define a previously unrecognized lineage of LepR-expressing CFs residing in the adventitia of coronary arteries and establish LepR-Cre as a tool to specifically label this CF subset.

### The LepR-Cre+ lineage arises and expands in neonatal stages

We next investigated the developmental timing of LepR-Cre+_CF emergence. E16.5 hearts were devoid of any LepR-Cre+ cells, but by postnatal day 1 (P1) a discontinuous layer of lineage-traced CFs appeared around developing arteries (Figure 2A). This lineage expanded rapidly after birth, and by P7 coronary arteries were fully encircled by LepR-Cre+_CFs, as seen in adulthood (Figure 2A). Unlike adult heart, which showed sparse labeling of interstitial CFs (Figure 1C,D), neonatal hearts contained no lineage-traced cells in interstitial regions. Flow cytometry confirmed that the LepR-Cre+_CF lineage expanded during the neonatal period, increasing from 2% of all ventricular fibroblasts at P1 to 14% by weaning (Figure 2B). Proliferation rates of LepR-Cre+ and LepR-Cre-CFs were comparable at P1 (25%), P7 (8%) and P19 (3%) (Figure 2C), suggesting that lineage expansion was driven by de novo *Lepr* expression rather than differential proliferation. RNAscope analyses supported this, revealing minimal *Lepr* expression at E16.5, but robust expression of both the long (*Leprb*) and short (*Lepra*) LepR isoforms in adventitial but not interstitial CFs or any other cardiac cell type at P1 and P7 (Figure 2D-G). <u>In the ventricular free walls, most CFs have an epicardial origin, but</u> in the septum a subset of CFs derives, during embryogenesis, from endocardial progenitors ^4^. <u>Analyses in *Tie2-Cre* lineage traced hearts revealed that, in regions containing endocardial-derived CFs, the pattern of *Lepr* expression was exactly the same as the one observed in the ventricular free walls: CFs occupying the adventitia of the septal coronary artery expressed *Lepr*, whereas interstitial CFs did not (</u>Supplemental Figure S1A-G)<u>. Together, these findings demonstrate that LepR-Cre+_CFs are not a distinct embryonic lineage, but instead arise during early neonatal life through de novo activation of *Lepr* in adventitial CFs of both epicardial– and endocardial-origin.</u>

**Figure 2.**
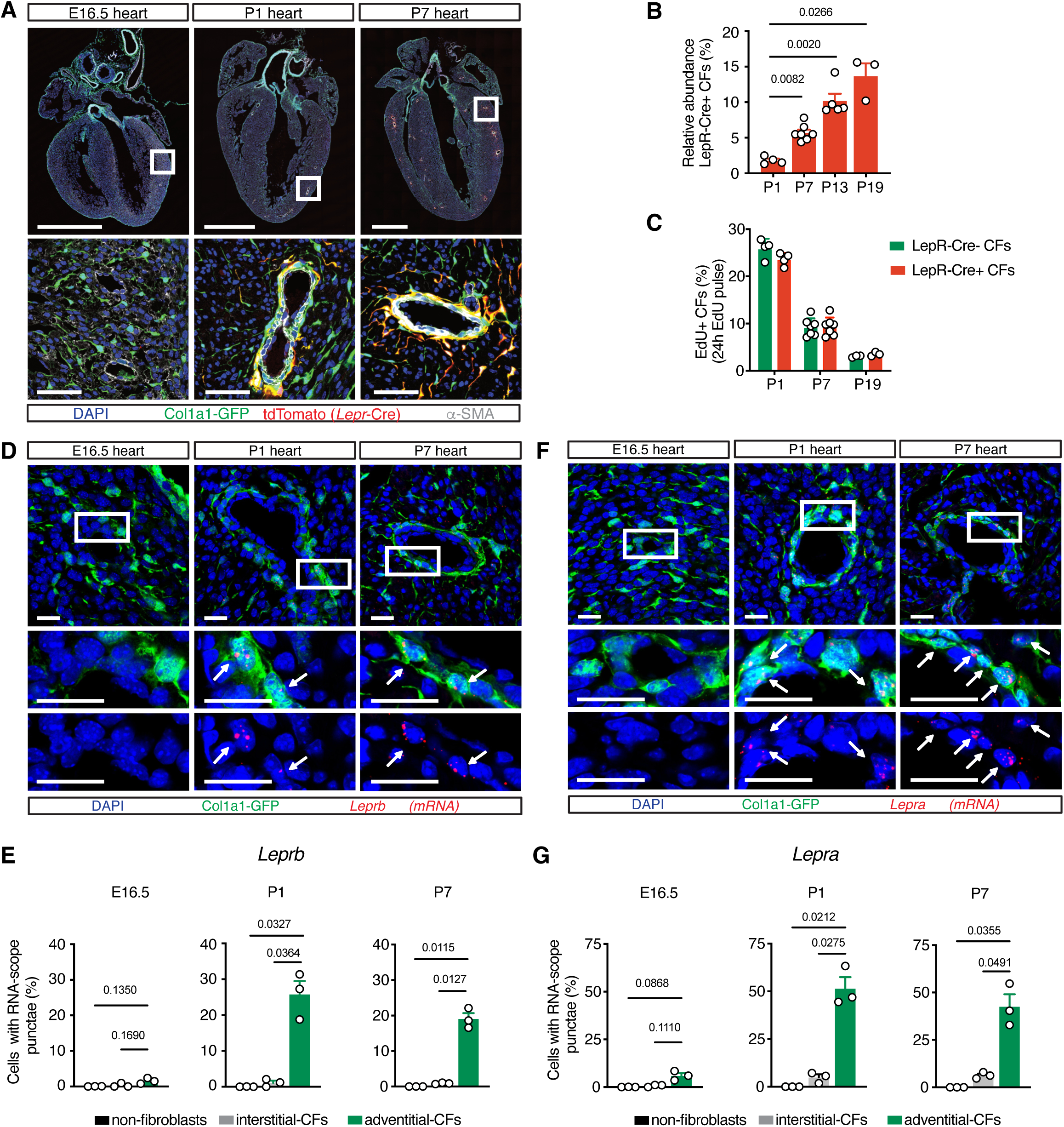
The LepR-Cre+ lineage of cardiac fibroblasts emerges and expands rapidly in the neonatal period. (**A**) Immunofluorescence time-course showing absence of Lepr-Cre+ cells in embryonic hearts (E16.5), appearance as a discontinuous monolayer around developing arteries at P1 and complete encirclement of coronary arteries with projections into adjacent tissue by P7 (*N* = 6). (**B**) Flow cytometry quantification showing progressive increase in the relative abundance of LepR-Cre+ fibroblasts from birth to weaning (*N* = 4 P1; 7 P7; 5 P13; 3 P19, mean ± SEM, one-way ANOVA). (**C**) EdU incorporation analysis revealing no significant difference in proliferative rates between LepR-Cre+ and LepR-Cre-neonatal fibroblasts (*N* = 4 P1; 7 P7; 3 P19, mean ± SEM, unpaired *t*-test). (**D-G**) RNA-scope analyses (D, F) and corresponding quantification (E, G) revealing expression of *Leprb* and *Lepra* (red dots) exclusively in adventitial fibroblasts (arrows; *N* = 3, mean ± SEM, one-way ANOVA). Boxed areas shown at higher magnification. Scale bars: 1 mm (whole-heart images), 50 µm (magnified panels).

### LepR+ cardiac fibroblasts express genes associated with progenitor potential

The neonatal activation of *Lepr* expression in adventitial CFs coincided with the onset of leptin production. At birth, mammals predominantly contain multilocular adipocytes with little to no leptin expression, as shown using a *Leptin-LacZ* reporter (Supplemental Figure S2). During the first postnatal week, subcutaneous adipose depots transition to a unilocular white phenotype and markedly increase leptin expression, leading to a surge in circulating leptin^26^. An ELISA time course (Figure 3A) confirmed that, in mice, circulating leptin was undetectable at birth, but rose sharply during the first neonatal week, peaking between P7 and P9 at values more than 4-fold higher than in lean adults (10 ng/ml). This temporal overlap between leptin production in fat and LepR expression in adventitial CFs suggested a previously unrecognized interorgan signaling axis between adipocytes and this CF subset.

**Figure 3.**
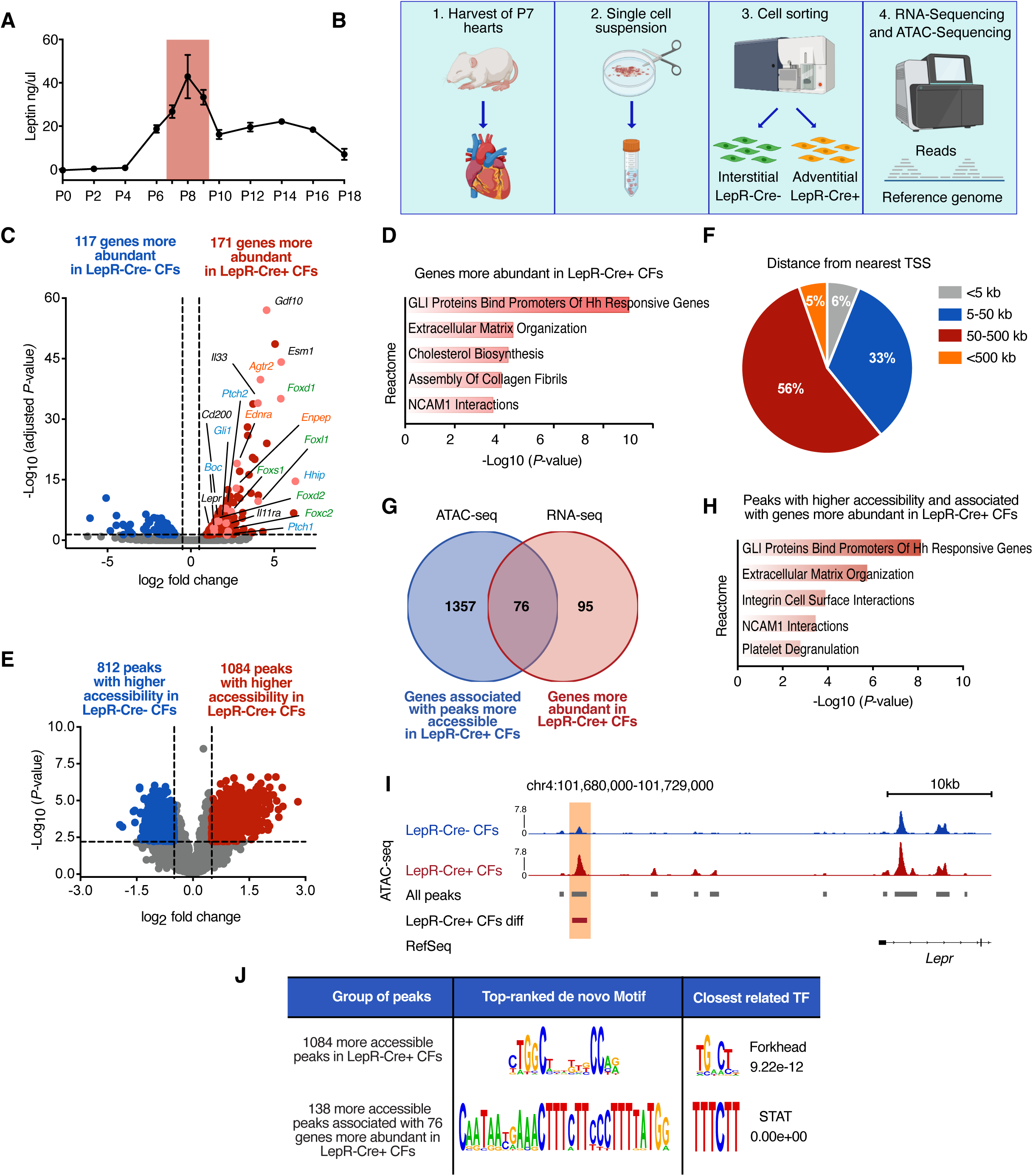
LepR-Cre+ cardiac fibroblasts display progenitor-associated transcriptional features and a distinct epigenetic landscape. (**A**) ELISA time-course showing a peak in circulating leptin between P7 and P9 (*N* = 3, mean ± SEM). (**B**) Experimental design for RNA-seq and ATAC-seq comparisons of neonatal adventitial LepR-Cre+ and interstitial LepR-Cre-fibroblasts. (**C**) Volcano plot showing differentially expressed genes (|log2FC|≥0.5, FDR ≤ 0.05, *N* = 3). Hedgehog pathway components (blue) and forkhead transcription factors (green) were enriched in LepR-Cre+_CFs. (**D**) Reactome analysis of genes with higher expression in LepR-Cre+ CFs showing enrichment of hedgehog signaling. (**E**) Volcano plot of ATAC-seq peaks with increased accessibility in LepR-Cre+_CFs (red) or LepR-Cre-_CFs (blue) (|log2FC|≥0.5, FDR ≤ 0.05, *N* = 2). (**F**) Distribution of ATAC-seq peaks relative to the nearest transcription start site (TSS). (**G**) Venn-diagram showing that almost half of the genes with higher expression in LepR-Cre+ CFs (RNA-seq) were associated with an ATAC-seq peak with higher accessibility in the same cells. (**H**) Reactome analysis of genes showing both higher expression in LepR-Cre+ CFs and associated with differential ATAC-seq peaks. (**I**) ATAC-seq tracks at the *Lepr* locus showing a peak (highlighted) with increased accessibility in LepR-Cre+ fibroblasts. (**J**) De novo motif analysis in differential ATAC-seq peaks. The top hit among all peaks more accessible in LepR-Cre+_CFs contained a Forkhead-like sequence, while a perfect STAT binding site was the top hit in the subset of peaks that were also associated with a gene showing increased expression in the same cells.

To explore molecular features distinguishing adventitial from interstitial CFs during the neonatal leptin peak, we performed RNA-seq on LepR-Cre+ and LepR-Cre-CFs FACS-sorted from P7 hearts (Figure 3B). A total of 288 differentially expressed genes (DEGs) were identified, with the strongest differences corresponding to genes expressed at higher levels in LepR-Cre+ than in LepR-Cre-fibroblasts, including *Lepr* itself (Figure 3C and Table S1). Reactome analysis of genes that had higher expression in LepR-Cre+_CFs (Figure 3D) revealed enrichment of transducers of hedgehog signaling, a pathway involved in the maintenance of tissue-resident stem cells and previously shown to be active in the neonatal coronary adventitial niche^27,28^. The most upregulated gene in LepR-Cre+_CFs was *Hhip* (6.27 logFC), a canonical hedgehog target and marker of pathway activation^29^. Also elevated were multiple forkhead transcription factors (*Foxd1*, *Foxl1*, *Foxd2*, *Foxc2*, *Foxs1*) known to work downstream of hedgehog signaling^30–32^. Additional top upregulated genes included *Gdf10* (a member of the BMP/TGFβ superfamily^33^) and *Esm1* (Endocan), an endothelial tip cell marker^34^ recently identified as a marker of bone marrow LepR+ mesenchymal stem cells^24^. Finally, LepR+_CFs displayed higher expression of IL33 and CD200, two key immunomodulatory molecules. Selected DEGs were validated via immunostaining and RNAscope (Supplemental Figure S3 A-C).

Collectively, these findings demonstrate that transcriptional differences between interstitial and adventitial CFs extend well beyond *Lepr*, with expression of stemness-associated genes being a distinguishing feature of adventitial fibroblasts.

### LepR-Cre+_CFs have a unique epigenetic landscape

To identify regulatory regions underlying gene expression differences between adventitial and interstitial cardiac fibroblasts, we performed ATAC-seq on LepR-Cre+ and LepR-Cre-CFs FACS-sorted from P7 hearts. From over 21,000 peaks of accessible chromatin (the majority farther than 50kb from the nearest TSS), 1084 were significantly more accessible in LepR-Cre_CFs (log₂FC ≥ 0.5, FDR ≤ 0.05, Figure 3E-F, Table S2). GREAT^35^ analysis linked these differential peaks to 1433 putative target genes (Table S2). Cross-referencing with RNA-seq results revealed that nearly half of the genes showing increased expression in LepR-Cre+_CFs were associated with at least one differential ATAC-seq peak (Figure 3G and Table S3), including key hedgehog pathway genes (*Ptch1, Ptch2*, *Boc*, *Gli1,* Figure 3H). We also identified a 1160bp region located 32kb upstream of the *Lepr* locus that may account for its selective expression in adventitial fibroblasts (Figure 3I). While most DEGs were associated with a single differential regulatory region (Supplemental Figure S4A), several of the genes showing strongest differential expression were associated with multiple differential elements – *Foxl1* (4.0 log2FC, 9 regions), *Foxd1* (5.4 log2FC, 5 regions) and *Gdf10* (4.5 log2FC, 4 regions) (Supplemental Figure S4B and Table S3). Motif analysis of the 1084 peaks enriched in LepR-Cre+_CFs identified a top de novo motif resembling a forkhead binding domain (Figure 3J). Restricting the analysis to peaks that had both greater accessibility in LepR-Cre+_CFs and were associated with genes expressed at higher levels in these cells revealed a top motif that contained a perfect match to the sequence recognized by STAT transcription factors, known nuclear mediators of leptin-LepR signaling^11,12^.

Together, these findings highlight specific regulatory elements that distinguish adventitial from interstitial CFs and suggest a model in which forkhead transcription factors promote the opening of specific regulatory regions and subsequent binding of STAT factors (elicited by leptin/LepR or other transduction pathways) to promote upregulation of target genes.

### Leptin signaling in LepR-Cre+_CFs

In addition to expressing *Lepr* transcripts, adventitial CFs also exhibited robust LepR protein levels (Supplemental Figure S5), suggesting they might be responsive to leptin. To test this, we FACS-sorted both LepR-Cre+_CFs and LepR-Cre-_CFs and stimulated them with leptin in vitro (Figure 4A). Western blot and immunocytochemistry revealed leptin-induced phosphorylation and nuclear accumulation of STAT3 exclusively in LepR-Cre+_CFs, but not in their interstitial LepR-Cre-counterparts, confirming that adventitial CFs are leptin-responsive (Figure 4B-D). We next examined whether leptin influences the transcriptome of adventitial CFs by performing RNA-seq on LepR-Cre+_CFs FACS-sorted from wildtype animals or Leptin KO/KO littermates at P7 (Figure 4E). The leptin null background resulted in modulation of 284 genes in LepR-Cre+_CFs (107 Down, 177 Up, Figure 4F, Table S4). Downregulated transcripts were significantly enriched in genes related to TGFβ signaling, a major pro-fibrotic pathway (Figure 4F). Consistently, *Thbs1* and *Ccn2* (encoding CTGF) were among the most downregulated transcripts. Other downregulated processes included response to shear stress (transcription factors KLF2 and KLF4) and negative regulation of signaling by smoothened. Conversely, genes upregulated in LepR-Cre+_CFs from Leptin null mutants were associated with ATP synthesis and myogenic development. Importantly, expression of hedgehog pathway components and forkhead transcription factors – key features distinguishing LepR-Cre+_CFs from their interstitial counterparts – remained unchanged in Leptin mutants, indicating that leptin is not required for maintenance of these lineage-defining transcriptional programs. Collectively, our data show that in neonatal adventitial CFs, leptin restrains myoblastic transcriptional programs and augments TGF-β target-gene expression as well as transcription factors linked to shear-stress sensing.

**Figure 4.**
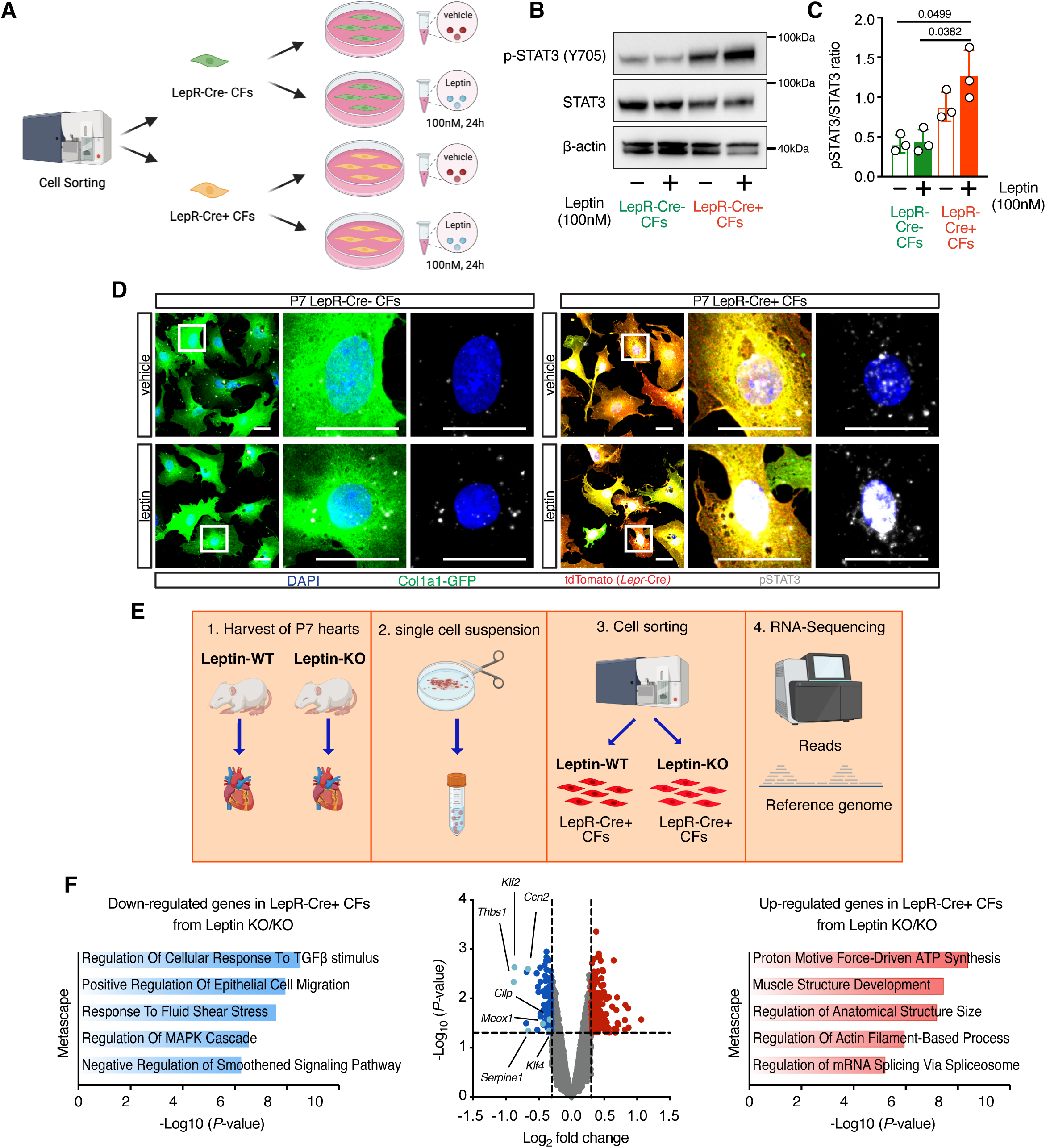
LepR-Cre+_CFs sense and respond to leptin. (**A**) Experimental design for ex vivo stimulation of CF subsets with leptin. (**B-C**) Western Blot (B) and respective quantification (C) showing increased STAT3 phosphorylation in LepR-Cre+_CFs after leptin stimulation (*N* = 3, mean ± SEM, one-way ANOVA). (**D**) Immunofluorescence showing nuclear accumulation of pSTAT3 (grey) specifically in LepR-Cre+ CFs after leptin stimulation (*N* = 3, scale bars = 25 µm). <u>(**E)** Experimental design to assess transcriptional alterations in LepR-Cre+_CFs isolated from leptin-deficient mice. (**F**) Volcano plot and functional annotation analyses of differentially expressed genes in LepR-Cre+_CFs from leptin mutants versus wild-type littermates (*N* = 4,</u> |log2FC|<u>≥ 0.3).</u>

### LepR-Cre+_CFs are preferential contributors to fibrosis

LepR+ mesenchymal cells have been extensively studied in bone marrow, where they act as tissue-resident progenitors that expand after injury to generate most newly formed bone^36,37^. Cardiac LepR-Cre+_CFs share features with these progenitors, including rapid neonatal expansion and expression of common markers (*Lepr*, *Esm1*)^24^, leading us to hypothesize that they might also function as progenitors during cardiac remodeling.

To test this, we subjected our triple transgenic animals to a surgical model of myocardial infarction (MI). Macroscopic analyses 28-day post-surgery revealed that, in contrast to the peri-vascular tdTomato pattern observed in sham hearts (arrows), in MI hearts lineage traced cells massively colonized the fibrotic scar (Figure 5A). <u>A histological time-course revealed that LepR-Cre+_CFs persisted in the infarcted area early post-MI (indicating that the ischemic shock does not trigger their death) and expanded rapidly after injury (</u>Figure 5B<u>). Quantification revealed that their relative abundance (LepR-Cre+_CFs as a fraction of all fibroblasts) doubled in the border zone within two days post-MI (</u>Figure 5C) and tripled in the scar one-week post-MI (Figure 5D). This significant increase in the relative abundance of LepR-Cre+_CFs suggested that, similarly to what happens in bone marrow^36^, in the heart LepR-Cre+ cells also expand preferentially to make a major contribution to tissue remodeling after injury. <u>Supporting this, EdU incorporation assays demonstrated higher proliferation rates in LepR-Cre+_CFs compared to their LepR-Cre-counterparts at 2-, 4– and 7-days post-MI (</u>Figure 5E, Supplemental Figure S6).

**Figure 5.**
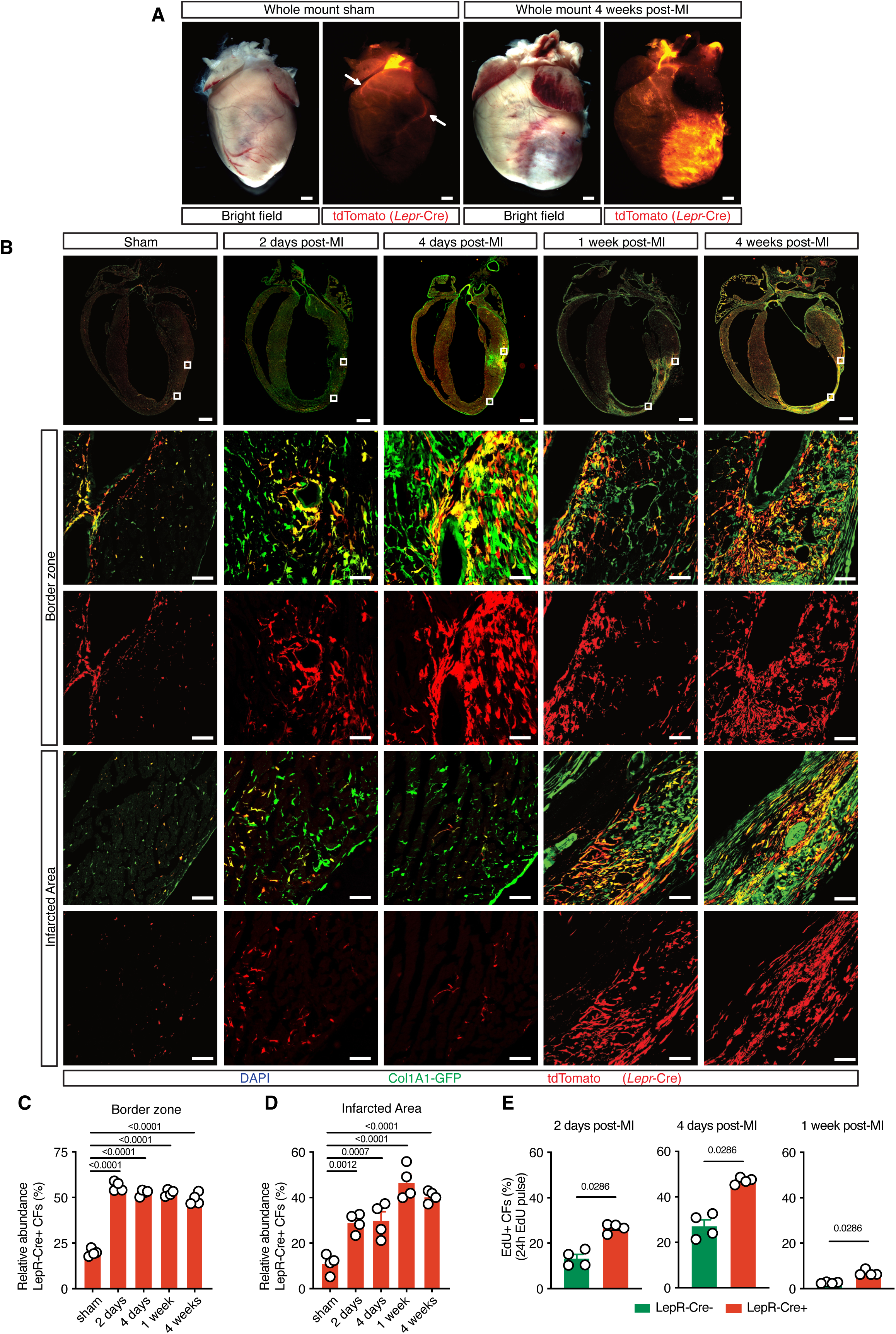
Adventitial LepR-Cre+ CFs are preferential contributors to fibrotic remodeling after MI. (**A**) Whole mount views of lineage-traced hearts showing the distribution of LepR-Cre+ CFs (tdTomato+). In sham hearts labeled cells were confined to the adventitia of large coronary arteries (arrows), but post-MI they colonized the entire scar. (**B**) Histological analysis of LepR-Cre+_CFs in distinct areas of the left ventricle before and after infarction. Boxed areas are shown at higher magnification. Scale bars: 1 mm (whole-heart images) and 100 µm (magnified panels). (**C-D**) <u>Quantification of histological data presented in (B), showing significant increases in the relative abundance of LepR-Cre+_CFs post-MI in the border zone (C) and scar (D) (*N* = 4, mean ± SEM,</u> one-way ANOVA<u>). (**E**) Quantitative analyses of EdU incorporation (histology in Figure S6) revealed higher proliferation rates in LepR-Cre+_CFs compared to LepR-Cre-counterparts at all time points post-MI (*N* = 4, mean ± SEM</u>, <u>Mann-Whitney test).</u>

The higher mitotic rates of LepR-Cre+_CFs explain the increase in their relative abundance post-injury, yet, because the LepR-Cre is constitutively active, lineage expansion could also reflect de novo *Lepr* activation post-injury. However, this possibility was refuted by analyses of published RNA-seq datasets^25,38^ and RNAscope demonstrating that *Lepr* expression declines after MI, with transcripts undetectable in scar fibroblasts (Supplemental Figure S7). These findings indicate that post-injury expansion of the LepR-Cre+ lineage reflects proliferation of pre-existing LepR-Cre+_CFs rather than new *Lepr* activation. To directly address this, we performed time-controlled lineage tracing using a tamoxifen-inducible LepR-CreERT2^24^. Neonatal tamoxifen induction of triple transgenic animals (*Lepr-CreERT2+; Col1a1-GFP+; Rosa-tdTomato/WT*) efficiently labeled adventitial CFs (Figure 6A-B, Supplemental Figure S8A), whereas adult induction produced weak labeling (Supplemental Figure S8B), consistent with lower adult *Lepr* expression observed via RNAscope (Figure 2 versus Supplemental Figure S7B,C). LepR-CreERT2 also labeled a small subset of cardiomyocytes (particularly evident in the atria), likely due to its lack of isoform specificity, unlike the constitutive LepR-Cre (Supplemental Figure S8A). When neonatal-induced animals were subjected to MI eight weeks later (Figure 6A), no new labeling could occur (Supplemental Figure S8C), yet the relative abundance of LepR-CreERT2+_CFs (LepR-CreERT2+_CFs as a fraction of all fibroblasts) still increased post-injury, both in the border zone and scar, to a similar extent as observed with the constitutive LepR-Cre (Figure 6C–E). These results confirm that pre-existing adventitial LepR-Cre+_CFs are preferential contributors to fibrotic remodeling following MI.

**Figure 6.**
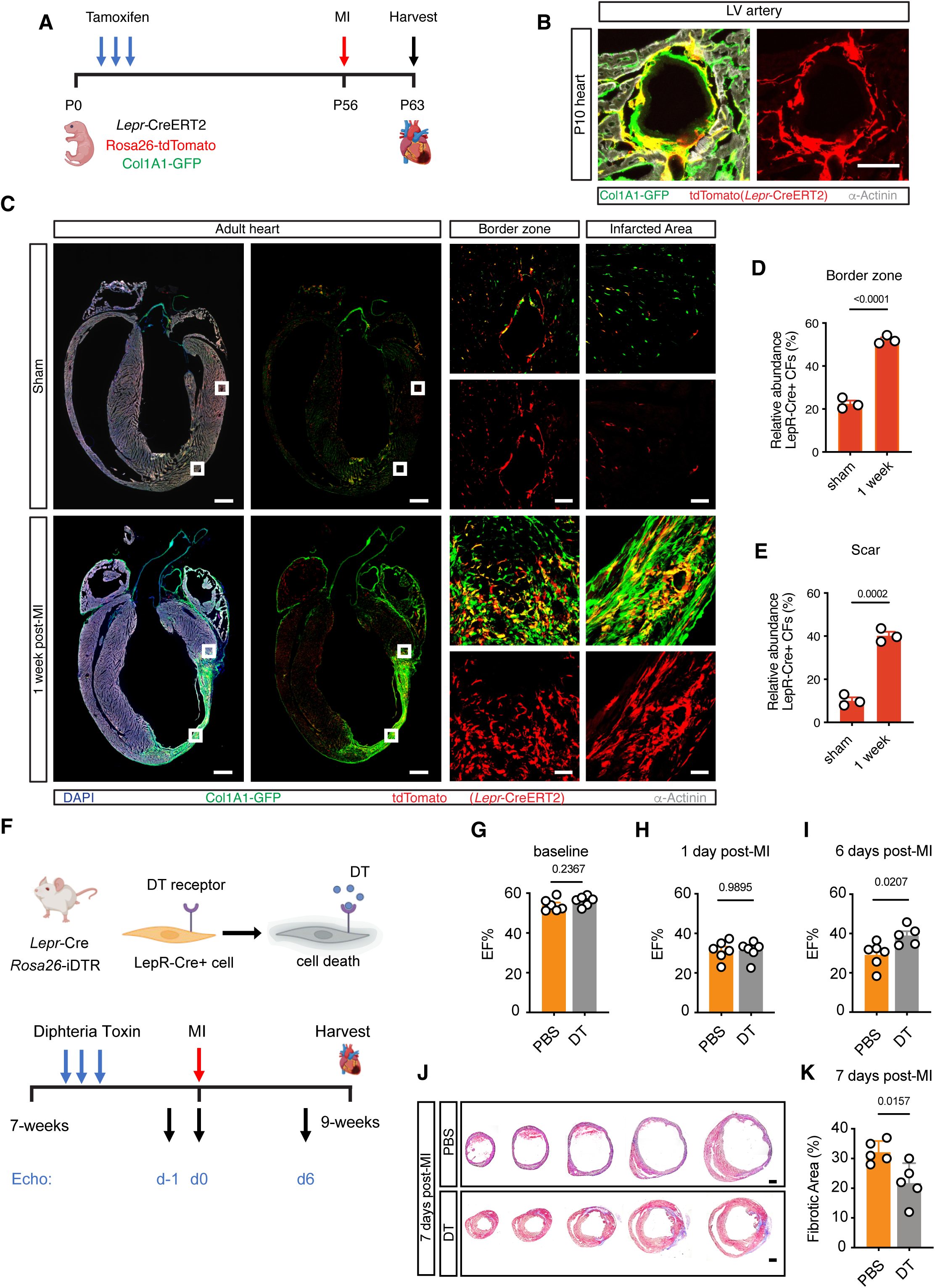
LepR-CreERT2+ fibroblasts expand post-MI, and depletion of LepR-Cre+fibroblasts improves outcomes post-MI. (**A**) Experimental timeline for time-controlled lineage tracing. LepR-CreERT2 was induced neonatally (P1-P3) and MI was induced 8 weeks later. (**B**) Immunofluorescence at P10 showing robust labeling of adventitial CFs by LepR-CreERT2 (*N* = 3, scale bar = 50µm). (**C**) Histological analysis of LepR-CreERT2+_CFs in different regions of the left ventricle pre– and post-infarction. Boxed areas are shown at higher magnification. Scale bars: 1mm (whole-heart images) and 100 µm (magnified panels). (**D-E**) Quantification of data in (C), showing a significant increase in the relative abundance of LepR-CreERT2+_CFs post-MI, both in the border zone (D) and scar (E) (*N* = 3, mean ± SEM, unpaired *t*-test). (**F**) Schematic of the strategy for LepR-Cre+_CF ablation. Adult *Lepr*-*Cre; Rosa26^iDTR^* mice were injected with PBS or diphtheria toxin (DT) for three consecutive days and subjected to MI five days later. (**G-I**) Echocardiographic assessment of left ventricular function showing improved ejection fraction (EF) 6 days after MI in DT-treated animals (*N* = 5-6, mean ± SEM, unpaired *t*-test). (**J,K**) Representative Masson’s trichrome-stained sections of infarcted hearts (J, scale bar = 1mm) and corresponding quantification showing reduced fibrotic area after LepR-Cre+_CF depletion (K, *N* = 5, mean ± SEM, unpaired *t*-test).

Finally, to test whether this preferential contribution extends beyond MI, we subjected LepR-Cre lineage traced mice to trans-aortic constriction (TAC), a model of cardiac hypertrophy and fibrosis without cardiomyocyte death^4^. As in MI, adventitial LepR-Cre+_CFs were also preferential contributors to fibrotic remodeling post-TAC (Supplemental Figure S9).

Collectively, these findings establish that the pre-existing lineage of LepR-Cre+ adventitial fibroblasts preferentially expands after cardiac injury and acts as a major progenitor pool driving fibrotic remodeling across distinct pathologic contexts.

### LepR-Cre+_CFs versus other fibroblast progenitors

Previous studies have described additional fibroblast progenitor lineages in the heart, including PDGFRα+/Sca1+ cells^39^, PW1+ cells^40^, HIC1+ cells^41^ and Gli1+ cells ^42^. Neither Sca1, *Peg3* (encoding PW1), nor *Hic1* were differently expressed between adventitial LepR-Cre+_CFs and their interstitial counterparts, indicating that the LepR-Cre+ lineage is distinct from these populations. However, the strong enrichment of Hedgehog pathway components in LepR-Cre+_CFs raised the possibility that they might overlap with Gli1-CreERT2+ cells. To test this, we compared LepR-Cre and Gli1-CreERT2 labeling profiles across multiple organs and observed strikingly different patterns (Supplemental Figures S10-S11). In the heart, Gli1-CreERT2, previously reported to label perivascular cells^42^, primarily marked fibroblasts. Unlike LepR-Cre, that was largely confined to the adventitia of large arteries, Gli1-CreERT2 labeled fibroblasts throughout the heart, including adventitial and interstitial fibroblasts in the ventricles, as well as fibroblasts in the atria and valves (Supplemental Figure S10). These results demonstrate that the two lineages are distinct and suggest LepR-Cre+_CFs represent a well-defined subpopulation within the broader pool of CFs labeled by Gli1-CreERT2.

### Improved post-MI outcomes upon depletion of LepR-Cre+_CFs

Our lineage tracing experiments identified LepR-Cre+_CFs to be preferentially expanded during fibrotic remodeling of the heart. To test the functional relevance of this population, we used *LepR-Cre; Rosa26-iDTR/WT* mice to ablate LepR-Cre+_CFs prior to surgical induction of MI. In these animals, cells with a history of *Lepr* expression become positive for the diphteria toxin receptor (normally not present in murine cells), rendering them susceptible to this toxin^23^, that was administered prior to MI or sham surgeries (Figure 6F). Echocardiographic analyses revealed no baseline differences between groups (Figure 6G and Supplemental Figure S12). The day after surgery, both genotype groups exhibited similar average reductions in ejection fraction (Figure 6H), revealing comparable injury size. However, by one-week post-MI, animals lacking LepR-Cre+_CFs displayed significantly better ejection fraction than controls without cell ablation (38.99% versus 29.22%, Figure 6I). Histological analyses further demonstrated that ablation of LepR-Cre⁺ CFs reduced scar size, as reflected by a decreased percentage of fibrotic area (21.8% of left ventricle area in animals with LepR-Cre+_CF depletion versus 32.2% in controls, Figure 6J,K).

These results demonstrate that LepR-Cre+_CFs play a key role in adverse remodeling after MI,and their depletion confers both structural and functional benefits.

### Transcriptional and epigenetic features distinguishing adult LepR-Cre+_CFs

To investigate transcriptional features underlying the preferential contribution of LepR-Cre+_CFs to the post-MI scar, we performed RNA-seq on LepR-Cre+ and LepR-Cre-CFs FACS-sorted from baseline adult hearts (P56, Figure 7A). Gene expression differences between adult CF subsets were less pronounced than at P7 (106 DEGs at P56 versus 288 DEGs at P7, Figures 7B and 3C, Table S5). Functional annotation of DEGs suggested that adventitial LepR-Cre+_CFs retained a more naïve transcriptome, consistent with a progenitor-like state (Figure 7C,D). Interstitial CFs (LepR-Cre-) expressed higher levels of WNT inhibitors, including *Wif1* and *Dkk3*, two genes known to mark CFs in an intermediate activation state but absent from CF progenitors^43^ (Figure 7B,C). Interstitial CFs also showed elevated expression of Scleraxis (*Scx*), a transcription factor driving CF activation^44^, as well as *Comp* and *Acta2*, two TGF-β1-inducible genes linked to fibroblast activation^45^. These patterns support that interstitial CFs are more activated than adventitial LepR-Cre+_CFs. Conversely, adventitial LepR-Cre+_CFs retained higher expression of molecules involved in hedgehog signal transduction, as previously observed at P7 (Figure 7B,D). However, transcriptional targets indicative of active hedgehog signaling (*Hhip* and most of the forkhead transcription factors) were no longer differentially expressed in adulthood, revealing reduced adventitial hedgehog activity with age, consistent with previous reports^28^. Nevertheless, adult adventitial LepR-Cre+_CFs retained all the machinery necessary to sense hedgehog ligands, positioning them in a privileged position to respond to increased hedgehog ligand availability in the heart post-MI^46^. Intersection analyses (Figure 7E,F) identified a core set of genes consistently enriched in LepR-Cre+_CFs at both P7 and P56 including *Lepr,* hedgehog-sensing components, *Esm1*, *Gdf10* and *Il33*. To complement transcriptomic data, we performed ATAC-on FACS-sorted adult CFs (Table S6). Joint analyses with neonatal datasets revealed large changes in chromatin accessibility between P7 and adulthood, with almost 9,000 differential peaks between these two stages in LepR-Cre+_CFs (Figure 7G and Table S6). As with transcriptomics analyses, chromatin differences between adventitial (LepR-Cre+) and interstitial CFs (LepR-Cre-) were less pronounced in adulthood. Multiple peaks that were significantly more accessible in LepR-Cre+_CFs at P7 trended toward higher accessibility in adulthood, but did not reach statistical thresholds for significance (Supplemental Figure S13).

**Figure 7.**
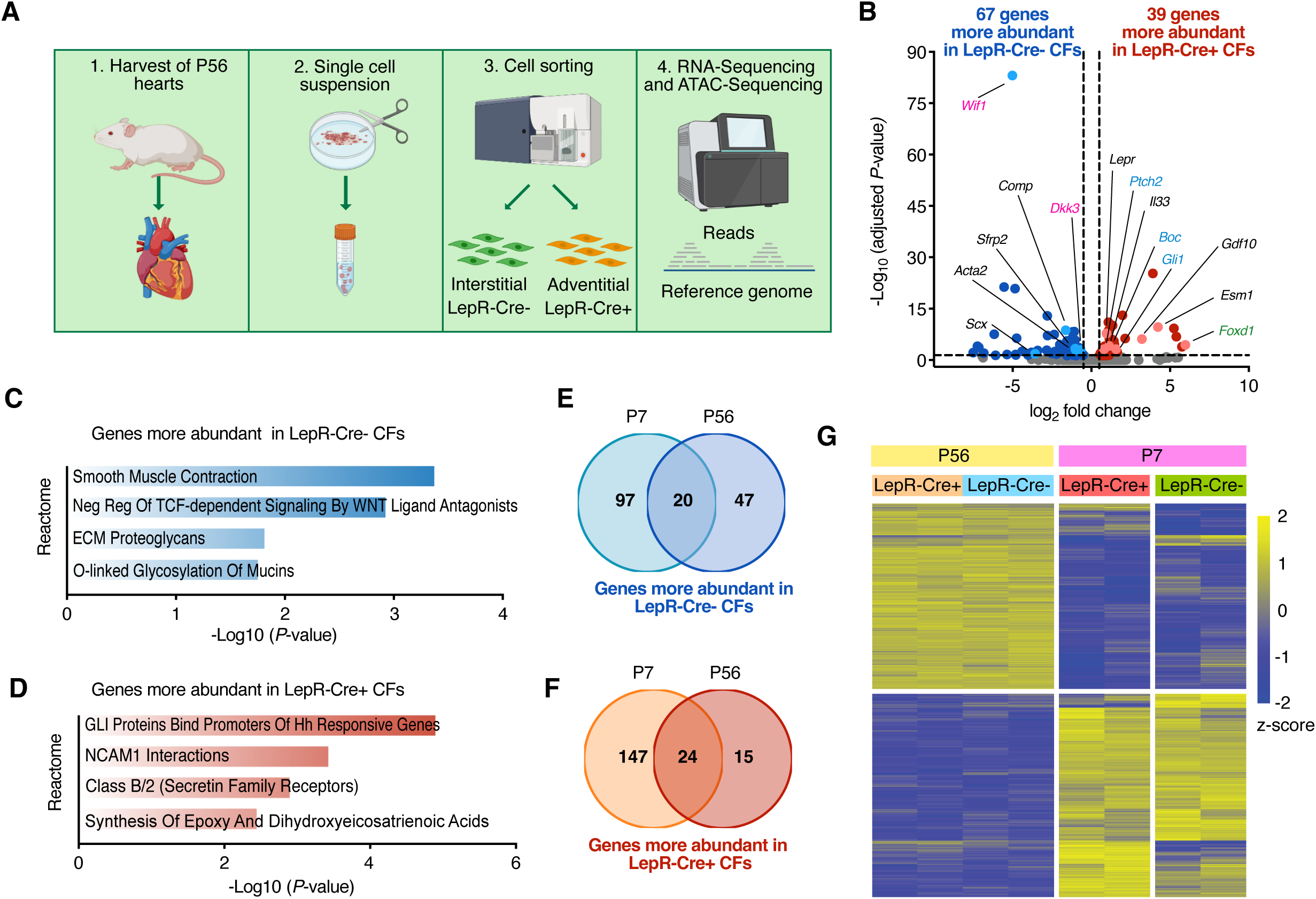
Transcriptomic and epigenomic profiling of adult LepR-Cre+ fibroblasts. (**A**) Schematic of the multiomics comparison between adult LepR-Cre+ and LepR-Cre-fibroblasts. (**B**) Volcano plot displaying transcripts significantly modulated (|log2FC|≥0.5, FDR ≤ 0.05) in LepR-Cre+ versus LepR-Cre-fibroblasts (*N* = 3). As in Figure 3D, hedgehog signaling molecules are highlighted in blue and forkhead transcription factors in green. (**C, D**) Reactome analysis of differentially expressed genes showing that LepR-Cre+_CFs had reduced expression of fibroblast activation markers (C) and increased expression of stemness-associated molecules (D), consistent with a progenitor-like state. (**E,F**) Venn diagrams showing genes consistently downregulated (E) or upregulated (F) in LepR-Cre+_CFs at both P7 and P56. (**G**) Heatmap of ATAC-seq data showing that cardiac maturation is associated with large-scale changes in fibroblast chromatin accessibility. Strong differences between LepR-Cre+ and LepR-Cre-fibroblasts at P7 were markedly attenuated in adulthood (*N* = 2).

Together, transcriptomic and epigenetic analyses reveal that adult adventitial LepR-Cre+_CFs were in a less activated state than interstitial CFs, maintaining progenitor-like features that may underlie their preferential contribution to cardiac remodeling.

### The heart as a potential source of leptin

In the regeneration-competent adult zebrafish heart, one of the two leptin paralogs is the most upregulated gene after injury ^47^. In humans MI triggers a peak in circulating leptin^48^, and previous reports suggested that cardiomyocytes in mice^49^ and rat^50,51^ may upregulate leptin expression after injury. These observations raised the possibility that, following MI, LepR-Cre+_CFs could sense leptin produced by neighboring cardiomyocytes.

To directly test for cardiac leptin production, we subjected *Leptin-LacZ/WT* mice to sham or MI surgery. X-Gal staining revealed no leptin expression in any cardiac cell type at baseline (Figure 8A). The only detectable signal was in brown/beige adipocytes surrounding the aortic and pulmonary arteries, which was markedly weaker than in visceral white adipocytes (Figure 8A). Importantly, this pattern remained remained unchanged after MI, with no evidence of leptin expression in cardiomyocytes or other cardiac cells at either 1-or 4-weeks post-injury (Figure 8A). Analysis of published RNA-seq datasets on purified cardiac cell types^25,38^ confirmed these findings. These results revealed that the peak of circulating leptin post-MI is not caused by production of this hormone in the heart, revealing differences between murine models and the regeneration-competent zebrafish heart.

**Figure 8.**
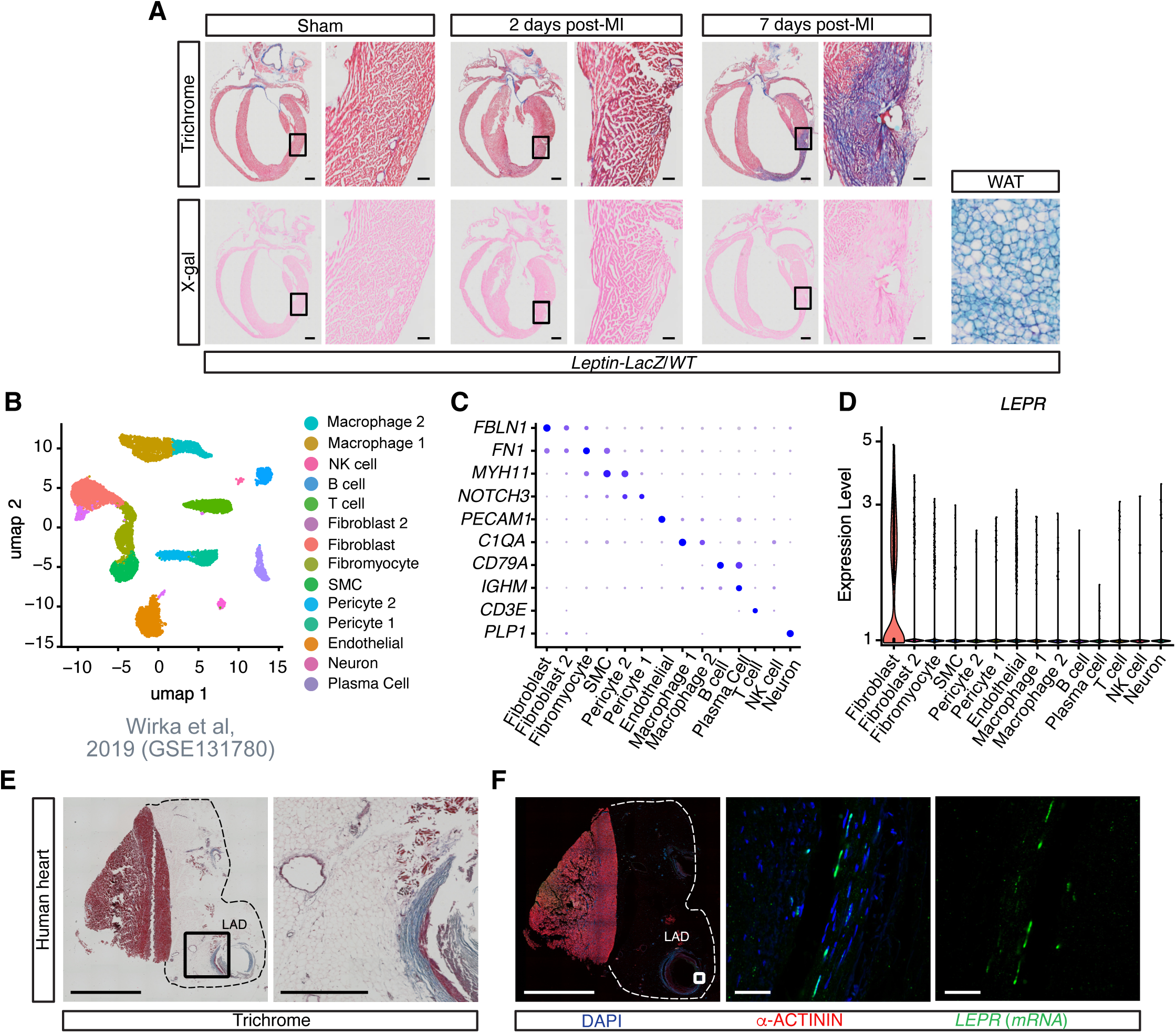
Adventitial fibroblasts in adipocyte-rich coronary niches of the human heart express *LEPR*. (**A**) Trichrome and X-gal staining of tissues from *Leptin-LacZ/WT* mice. No X-gal-positive cells were detected in the heart before or after MI. Peri-gonadal white adipose tissue (WAT) served as a positive control. Boxed areas are shown at higher magnification (*N* = 3, scale bars: 1 mm for whole-heart images, 100 µm for magnified panels). <u>(**B**) UMAP plot showing cell types/clusters in scRNA-seq of cells isolated from the human right coronary artery (*N* = 4). (**C**) Dot plot showing expression of top cluster markers. (**D**) *LEPR* expression across clusters, highlighting adventitial fibroblasts as the main expressing population. (**E,F**) Trichrome (E) and RNAscope combined with immunofluorescence (F) of human heart sections showing an LAD imbedded in abundant adipocytes (E), with *LEPR* expression (green) detected in adventitial fibroblasts (F). Dashed lines in (E,F) delineate the borders of the epicardial adipose tissue. Boxed areas are shown at higher magnification (*N* = 3, scale bars: 5 mm for whole-heart images, 100 µm for magnified panels).</u>

Unlike the largely fat-free mouse heart, the human heart contains abundant adipocytes surrounding large coronary arteries (Figure 8E). Thus, if adventitial LepR+_CFs also exist in humans, they would be directly embedded in an adipokine-rich environment. To investigate this, we analyzed published scRNA-seq datasets of the human heart. In datasets from interstitial myocardium (areas devoid of large coronary arteries), *LEPR* was absent from CFs, but, depending on the source laboratory, was detectable in endocardial cells ^52–54^(Supplemental Figure S14). In contrast, re-analysis of a dataset specifically probing coronary arteries^55^ allowed us to identify *LEPR* as a defining feature of adventitial fibroblasts, mirroring our murine findings (Figure 8B-D). To further validate this observation, we performed RNAscope analyses of human heart sections containing large segments of the LAD, and confirmed abundant expression of the long *LEPR* isoform specifically in adventitial cells (Figure 8E,F).

Together, these findings demonstrate evolutionary conservation of adventitial LepR+CFs in mammals, refute the idea of cardiomyocytes as a cardiac leptin source, and suggest that adipocytes – rare in mice but abundant in humans – are the sole producers of leptin in the mammalian heart.

## DISCUSSION

Understanding cellular and molecular mechanisms underlying cardiac fibrosis, a hallmark of most forms of heart failure, is of paramount importance. Here, using state-of-the-art approaches for gene expression profiling, imaging and lineage-tracing, we identify a unique population of CFs localized to the adventitia of large coronary arteries and characterized by LepR expression that serve as preferential contributors to fibrotic remodeling of the heart. <u>Importantly, we also identified LepR+ adventitial CFs in the human heart. Notably, in mice, depletion of LepR+ CFs resulted in improved outcomes post-MI.</u>

In addition to *Lepr*, a defining feature distinguishing adventitial CFs from their interstitial counterparts is expression of mediators of hedgehog signaling, a pathway implicated in the maintenance of tissue-resident progenitors^27^. <u>These hedgehog-related genes were not modulated in LepR-Cre+_CFs from leptin-deficient mice, indicating that their expression is not regulated by leptin</u>. Although transcriptomic and epigenetic differences between CF subsets diminish with age, adult LepR-Cre+_CFs retain the machinery required to sense hedgehog ligands, which are strongly induced after MI^46^. <u>This feature may underlie their preferential contribution to fibrotic remodeling.</u>

The existence of a LepR-Cre+ population of mesenchymal progenitors in the heart is a novel finding, though an analogous population has been recognized in bone marrow^36^. Strikingly, in addition to sharing markers (*Lepr, Esm1*), these populations also share developmental timing in the neonatal period^24,37^, coinciding with onset of leptin production. This parallel suggests that LepR+ mesenchymal progenitors may emerge <u>simultaneously across multiple tissues, raising the possibility that similar mesenchymal populations exist in additional organs, as supported by our multi-tissue analyses.</u> <u>It would be of interest to test whether LepR-Cre+ fibroblasts are preferential contributors to fibrotic remodeling in other organs where *Lepr* expression is fibroblast-restricted, such as the lung.</u> In addition to their role in post-injury repair, bone marrow LepR-Cre+ cells also maintain the hematopoietic niche through secretion of trophic factors. Our finding that cardiac LepR-Cre+_CFs express signaling molecules such as IL33 and GDF10 raises the possibility of a similar niche-maintaining role.

Our findings also compel a reappraisal of the role of leptin in cardiac biology. Leptin has been implicated in hypertrophy, which could be a primary effect or an indirect consequence elicited by changes in blood pressure (another known action of leptin)^17^. Prior studies, many relying on limited technologies, proposed that leptin acts directly on cardiomyocytes^17^. However, our data – spanning RNA-seq, RNAscope and genetic lineage tracing – demonstrated that, in the murine heart, CFs are the only cell type expressing the signaling-capable long isoform of LepR. This is seemingly in contrast with previous cardiomyocyte-centered reports^13–16^, but it is possible that alternative receptors or shorter LepR isoforms with unrecognized signaling capability might be involved. Furthermore, contrary to earlier reports^49–51^, we found no evidence for cardiac leptin production in mice either at baseline or post-MI. This is in contrast to the regeneration-competent adult zebrafish heart, where leptin is one of the top upregulated genes post-injury^47^. Given that pro-fibrotic cues in mammals can promote regenerative responses in zebrafish^45,56^, investigating how leptin is sensed in the zebrafish heart could yield insight into species-specific regenerative mechanisms.

In conclusion, we identified a new population of adventitial LepR-Cre+ fibroblasts that act as progenitors driving fibrotic remodeling of the heart. Beyond reshaping our understanding of processes underlying cardiac fibrosis, our findings also revise the paradigm of leptin/LepR signaling in the heart and open new avenues for exploring how this major endocrine pathway intersects with cardiac disease.

## ACKNOWLEDGMENTS and SOURCES OF FUNDING

This work was supported by funds from the German Center for Cardiovascular Research (DZHK, Junior Group Leader grant to NG-C), Italian Ministry of Health (Ricerca Corrente to Centro Cardiologico Monzino IRCCS), Leducq foundation (NG-C and SME), Cardio Pulmonary Institute (PC), Johanna Quandt Stiftung (NG-C) and Giuliani Foundation (NG-C). VL received a postdoctoral fellowship from the Humboldt foundation and OA received a fellowship from the IEO-Monzino Foundation.

## REFERENCES

1. Roth GA, Mensah GA, Johnson CO, Addolorato G, Ammirati E, Baddour LM, Barengo NC, Beaton AZ, Benjamin EJ, Benziger CP, et al. Global Burden of Cardiovascular Diseases and Risk Factors, 1990-2019: Update From the GBD 2019 Study. J Am Coll Cardiol. 2020;76:2982–3021. doi: 10.1016/j.jacc.2020.11.010

2. Tallquist MD, Molkentin JD. Redefining the identity of cardiac fibroblasts. Nat Rev Cardiol. 2017;14:484–491. doi: 10.1038/nrcardio.2017.57

3. Frangogiannis NG. Cardiac fibrosis. Cardiovasc Res. 2021;117:1450–1488. doi: 10.1093/cvr/cvaa324

4. Moore-Morris T, Guimarães-Camboa N, Banerjee I, Zambon AC, Kisseleva T, Velayoudon A, Stallcup WB, Gu Y, Dalton ND, Cedenilla M, et al. Resident fibroblast lineages mediate pressure overload-induced cardiac fibrosis. J Clin Invest. 2014;124:2921–2934. doi: 10.1172/JCI74783

5. Moore-Morris T, Cattaneo P, Guimarães-Camboa N, Bogomolovas J, Cedenilla M, Banerjee I, Ricote M, Kisseleva T, Zhang L, Gu Y, et al. Infarct Fibroblasts Do Not Derive From Bone Marrow Lineages. Circ Res. 2018;122:583–590. doi: 10.1161/CIRCRESAHA.117.311490

6. Ali SR, Ranjbarvaziri S, Talkhabi M, Zhao P, Subat A, Hojjat A, Kamran P, Müller AM, Volz KS, Tang Z, et al. Developmental heterogeneity of cardiac fibroblasts does not predict pathological proliferation and activation. Circ Res. 2014;115:625–635. doi: 10.1161/CIRCRESAHA.115.303794

7. Fu X, Khalil H, Kanisicak O, Boyer JG, Vagnozzi RJ, Maliken BD, Sargent MA, Prasad V, Valiente-Alandi I, Blaxall BC, et al. Specialized fibroblast differentiated states underlie scar formation in the infarcted mouse heart. J Clin Invest. 2018;128:2127–2143. doi: 10.1172/JCI98215

8. Aghajanian H, Kimura T, Rurik JG, Hancock AS, Leibowitz MS, Li L, Scholler J, Monslow J, Lo A, Han W, et al. Targeting cardiac fibrosis with engineered T cells. Nature. 2019;573:430–433. doi: 10.1038/s41586-019-1546-z

9. Kenchaiah S, Evans JC, Levy D, Wilson PW, Benjamin EJ, Larson MG, Kannel WB, Vasan RS. Obesity and the risk of heart failure. N Engl J Med. 2002;347:305–313. doi: 10.1056/NEJMoa020245

10. Scheja L, Heeren J. The endocrine function of adipose tissues in health and cardiometabolic disease. Nat Rev Endocrinol. 2019;15:507–524. doi: 10.1038/s41574-019-0230-6

11. Friedman JM. Leptin and the endocrine control of energy balance. Nat Metab. 2019;1:754–764. doi: 10.1038/s42255-019-0095-y

12. Peelman F, Zabeau L, Moharana K, Savvides SN, Tavernier J. 20 years of leptin: insights into signaling assemblies of the leptin receptor. J Endocrinol. 2014;223:T9–23. doi: 10.1530/JOE-14-0264

13. Rajapurohitam V, Gan XT, Kirshenbaum LA, Karmazyn M. The obesity-associated peptide leptin induces hypertrophy in neonatal rat ventricular myocytes. Circ Res. 2003;93:277–279. doi: 10.1161/01.RES.0000089255.37804.72

14. Madani S, De Girolamo S, Muñoz DM, Li RK, Sweeney G. Direct effects of leptin on size and extracellular matrix components of human pediatric ventricular myocytes. Cardiovasc Res. 2006;69:716–725. doi: 10.1016/j.cardiores.2005.11.022

15. McGaffin KR, Witham WG, Yester KA, Romano LC, O’Doherty RM, McTiernan CF, O’Donnell CP. Cardiac-specific leptin receptor deletion exacerbates ischaemic heart failure in mice. Cardiovasc Res. 2011;89:60–71. doi: 10.1093/cvr/cvq288

16. Hall ME, Smith G, Hall JE, Stec DE. Cardiomyocyte-specific deletion of leptin receptors causes lethal heart failure in Cre-recombinase-mediated cardiotoxicity. Am J Physiol Regul Integr Comp Physiol. 2012;303:R1241–1250. doi: 10.1152/ajpregu.00292.2012

17. Mao Y, Zhao K, Li P, Sheng Y. The emerging role of leptin in obesity-associated cardiac fibrosis: evidence and mechanism. Mol Cell Biochem. 2023;478:991–1011. doi: 10.1007/s11010-022-04562-6

18. DeFalco J, Tomishima M, Liu H, Zhao C, Cai X, Marth JD, Enquist L, Friedman JM. Virus-assisted mapping of neural inputs to a feeding center in the hypothalamus. Science. 2001;291:2608–2613. doi: 10.1126/science.1056602

19. Yata Y, Scanga A, Gillan A, Yang L, Reif S, Breindl M, Brenner DA, Rippe RA. DNase I-hypersensitive sites enhance alpha1(I) collagen gene expression in hepatic stellate cells. Hepatology. 2003;37:267–276. doi: 10.1053/jhep.2003.50067

20. Kisanuki YY, Hammer RE, Miyazaki J, Williams SC, Richardson JA, Yanagisawa M. Tie2-Cre transgenic mice: a new model for endothelial cell-lineage analysis in vivo. Dev Biol. 2001;230:230–242. doi: 10.1006/dbio.2000.0106

21. Ahn S, Joyner AL. Dynamic changes in the response of cells to positive hedgehog signaling during mouse limb patterning. Cell. 2004;118:505–516. doi: 10.1016/j.cell.2004.07.023

22. Madisen L, Zwingman TA, Sunkin SM, Oh SW, Zariwala HA, Gu H, Ng LL, Palmiter RD, Hawrylycz MJ, Jones AR, et al. A robust and high-throughput Cre reporting and characterization system for the whole mouse brain. Nat Neurosci. 2010;13:133–140. doi: 10.1038/nn.2467

23. Buch T, Heppner FL, Tertilt C, Heinen TJ, Kremer M, Wunderlich FT, Jung S, Waisman A. A Cre-inducible diphtheria toxin receptor mediates cell lineage ablation after toxin administration. Nat Methods. 2005;2:419–426. doi: 10.1038/nmeth762

24. Sivaraj KK, Jeong HW, Dharmalingam B, Zeuschner D, Adams S, Potente M, Adams RH. Regional specialization and fate specification of bone stromal cells in skeletal development. Cell Rep. 2021;36:109352. doi: 10.1016/j.celrep.2021.109352

25. Quaife-Ryan GA, Sim CB, Ziemann M, Kaspi A, Rafehi H, Ramialison M, El-Osta A, Hudson JE, Porrello ER. Multicellular Transcriptional Analysis of Mammalian Heart Regeneration. Circulation. 2017;136:1123–1139. doi: 10.1161/CIRCULATIONAHA.117.028252

26. Ahima RS, Prabakaran D, Flier JS. Postnatal leptin surge and regulation of circadian rhythm of leptin by feeding. Implications for energy homeostasis and neuroendocrine function. J Clin Invest. 1998;101:1020–1027. doi: 10.1172/JCI1176

27. Petrova R, Joyner AL. Roles for Hedgehog signaling in adult organ homeostasis and repair. Development. 2014;141:3445–3457. doi: 10.1242/dev.083691

28. Passman JN, Dong XR, Wu SP, Maguire CT, Hogan KA, Bautch VL, Majesky MW. A sonic hedgehog signaling domain in the arterial adventitia supports resident Sca1+ smooth muscle progenitor cells. Proc Natl Acad Sci U S A. 2008;105:9349–9354. doi: 10.1073/pnas.0711382105

29. Chuang PT, McMahon AP. Vertebrate Hedgehog signalling modulated by induction of a Hedgehog-binding protein. Nature. 1999;397:617–621. doi: 10.1038/17611

30. Fink DM, Sun MR, Heyne GW, Everson JL, Chung HM, Park S, Sheets MD, Lipinski RJ. Coordinated d-cyclin/Foxd1 activation drives mitogenic activity of the Sonic Hedgehog signaling pathway. Cell Signal. 2018;44:1–9. doi: 10.1016/j.cellsig.2017.12.007

31. Madison BB, McKenna LB, Dolson D, Epstein DJ, Kaestner KH. FoxF1 and FoxL1 link hedgehog signaling and the control of epithelial proliferation in the developing stomach and intestine. J Biol Chem. 2009;284:5936–5944. doi: 10.1074/jbc.M808103200

32. Rowton M, Perez-Cervantes C, Hur S, Jacobs-Li J, Lu E, Deng N, Guzzetta A, Hoffmann AD, Stocker M, Steimle JD, et al. Hedgehog signaling activates a mammalian heterochronic gene regulatory network controlling differentiation timing across lineages. Dev Cell. 2022;57:2181–2203.e2189. doi: 10.1016/j.devcel.2022.08.009

33. Upadhyay G, Yin Y, Yuan H, Li X, Derynck R, Glazer RI. Stem cell antigen-1 enhances tumorigenicity by disruption of growth differentiation factor-10 (GDF10)-dependent TGF-beta signaling. Proc Natl Acad Sci U S A. 2011;108:7820–7825. doi: 10.1073/pnas.1103441108

34. Rocha SF, Schiller M, Jing D, Li H, Butz S, Vestweber D, Biljes D, Drexler HC, Nieminen-Kelhä M, Vajkoczy P, et al. Esm1 modulates endothelial tip cell behavior and vascular permeability by enhancing VEGF bioavailability. Circ Res. 2014;115:581–590. doi: 10.1161/CIRCRESAHA.115.304718

35. McLean CY, Bristor D, Hiller M, Clarke SL, Schaar BT, Lowe CB, Wenger AM, Bejerano G. GREAT improves functional interpretation of cis-regulatory regions. Nat Biotechnol. 2010;28:495–501. doi: 10.1038/nbt.1630

36. Zhou BO, Yue R, Murphy MM, Peyer JG, Morrison SJ. Leptin-receptor-expressing mesenchymal stromal cells represent the main source of bone formed by adult bone marrow. Cell Stem Cell. 2014;15:154–168. doi: 10.1016/j.stem.2014.06.008

37. Shu HS, Liu YL, Tang XT, Zhang XS, Zhou B, Zou W, Zhou BO. Tracing the skeletal progenitor transition during postnatal bone formation. Cell Stem Cell. 2021;28:2122–2136.e2123. doi: 10.1016/j.stem.2021.08.010

38. Rogg EM, Abplanalp WT, Bischof C, John D, Schulz MH, Krishnan J, Fischer A, Poluzzi C, Schaefer L, Bonauer A, et al. Analysis of Cell Type-Specific Effects of MicroRNA-92a Provides Novel Insights Into Target Regulation and Mechanism of Action. Circulation. 2018;138:2545–2558. doi: 10.1161/CIRCULATIONAHA.118.034598

39. Chong JJ, Chandrakanthan V, Xaymardan M, Asli NS, Li J, Ahmed I, Heffernan C, Menon MK, Scarlett CJ, Rashidianfar A, et al. Adult cardiac-resident MSC-like stem cells with a proepicardial origin. Cell Stem Cell. 2011;9:527–540. doi: 10.1016/j.stem.2011.10.002

40. Yaniz-Galende E, Roux M, Nadaud S, Mougenot N, Bouvet M, Claude O, Lebreton G, Blanc C, Pinet F, Atassi F, et al. Fibrogenic Potential of PW1/Peg3 Expressing Cardiac Stem Cells. J Am Coll Cardiol. 2017;70:728–741. doi: 10.1016/j.jacc.2017.06.010

41. Soliman H, Paylor B, Scott RW, Lemos DR, Chang C, Arostegui M, Low M, Lee C, Fiore D, Braghetta P, et al. Pathogenic Potential of Hic1-Expressing Cardiac Stromal Progenitors. Cell Stem Cell. 2020;26:459–461. doi: 10.1016/j.stem.2020.01.023

42. Kramann R, Schneider RK, DiRocco DP, Machado F, Fleig S, Bondzie PA, Henderson JM, Ebert BL, Humphreys BD. Perivascular Gli1+ progenitors are key contributors to injury-induced organ fibrosis. Cell Stem Cell. 2015;16:51–66. doi: 10.1016/j.stem.2014.11.004

43. Farbehi N, Patrick R, Dorison A, Xaymardan M, Janbandhu V, Wystub-Lis K, Ho JW, Nordon RE, Harvey RP. Single-cell expression profiling reveals dynamic flux of cardiac stromal, vascular and immune cells in health and injury. Elife. 2019;8. doi: 10.7554/eLife.43882

44. Nagalingam RS, Chattopadhyaya S, Al-Hattab DS, Cheung DYC, Schwartz LY, Jana S, Aroutiounova N, Ledingham DA, Moffatt TL, Landry NM, et al. Scleraxis and fibrosis in the pressure-overloaded heart. Eur Heart J. 2022. doi: 10.1093/eurheartj/ehac362

45. Schafer S, Viswanathan S, Widjaja AA, Lim WW, Moreno-Moral A, DeLaughter DM, Ng B, Patone G, Chow K, Khin E, et al. IL-11 is a crucial determinant of cardiovascular fibrosis. Nature. 2017;552:110–115. doi: 10.1038/nature24676

46. Kusano KF, Pola R, Murayama T, Curry C, Kawamoto A, Iwakura A, Shintani S, Ii M, Asai J, Tkebuchava T, et al. Sonic hedgehog myocardial gene therapy: tissue repair through transient reconstitution of embryonic signaling. Nat Med. 2005;11:1197–1204. doi: 10.1038/nm1313

47. Kang J, Hu J, Karra R, Dickson AL, Tornini VA, Nachtrab G, Gemberling M, Goldman JA, Black BL, Poss KD. Modulation of tissue repair by regeneration enhancer elements. Nature. 2016;532:201–206. doi: 10.1038/nature17644

48. Khafaji HA, Bener AB, Rizk NM, Al Suwaidi J. Elevated serum leptin levels in patients with acute myocardial infarction; correlation with coronary angiographic and echocardiographic findings. BMC Res Notes. 2012;5:262. doi: 10.1186/1756-0500-5-262

49. Kain D, Simon AJ, Greenberg A, Ben Zvi D, Gilburd B, Schneiderman J. Cardiac leptin overexpression in the context of acute MI and reperfusion potentiates myocardial remodeling and left ventricular dysfunction. PLoS One. 2018;13:e0203902. doi: 10.1371/journal.pone.0203902

50. Purdham DM, Zou MX, Rajapurohitam V, Karmazyn M. Rat heart is a site of leptin production and action. Am J Physiol Heart Circ Physiol. 2004;287:H2877–2884. doi: 10.1152/ajpheart.00499.2004

51. Matsui H, Motooka M, Koike H, Inoue M, Iwasaki T, Suzuki T, Kurabayashi M, Yokoyama T. Ischemia/reperfusion in rat heart induces leptin and leptin receptor gene expression. Life Sci. 2007;80:672–680. doi: 10.1016/j.lfs.2006.10.027

52. Litviòuková M, Talavera-López C, Maatz H, Reichart D, Worth CL, Lindberg EL, Kanda M, Polanski K, Heinig M, Lee M, et al. Cells of the adult human heart. Nature. 2020;588:466–472. doi: 10.1038/s41586-020-2797-4

53. Nicin L, Schroeter SM, Glaser SF, Schulze-Brüning R, Pham MD, Hille SS, Yekelchyk M, Kattih B, Abplanalp WT, Tombor L, et al. A human cell atlas of the pressure-induced hypertrophic heart. Nat Cardiovasc Res. 2022;1:174–185. doi: 10.1038/s44161-022-00019-7

54. Koenig AL, Shchukina I, Amrute J, Andhey PS, Zaitsev K, Lai L, Bajpai G, Bredemeyer A, Smith G, Jones C, et al. Single-cell transcriptomics reveals cell-type-specific diversification in human heart failure. Nat Cardiovasc Res. 2022;1:263–280. doi: 10.1038/s44161-022-00028-6

55. Wirka RC, Wagh D, Paik DT, Pjanic M, Nguyen T, Miller CL, Kundu R, Nagao M, Coller J, Koyano TK, et al. Atheroprotective roles of smooth muscle cell phenotypic modulation and the TCF21 disease gene as revealed by single-cell analysis. Nat Med. 2019;25:1280–1289. doi: 10.1038/s41591-019-0512-5

56. Allanki S, Strilic B, Scheinberger L, Onderwater YL, Marks A, Günther S, Preussner J, Kikhi K, Looso M, Stainier DYR, et al. Interleukin-11 signaling promotes cellular reprogramming and limits fibrotic scarring during tissue regeneration. Sci Adv. 2021;7:eabg6497. doi: 10.1126/sciadv.abg6497

